# *In vivo* measurements reveal a single 5’ intron is sufficient to increase protein expression level in *C. elegans*

**DOI:** 10.1101/499459

**Authors:** Matthew M. Crane, Bryan Sands, Christian Battaglia, Brock Johnson, Soo Yun, Matt Kaeberlein, Roger Brent, Alex Mendenhall

## Abstract

Introns can increase gene expression levels using a variety of mechanisms collectively referred to as Intron Mediated Enhancement (IME). To date, the magnitude of IME has been quantified in human cell culture and plant models by comparing intronless reporter gene expression levels to those of intron-bearing reporter genes *in vitro* (mRNA, Western Blots, protein activity), using genome editing technologies that lacked full control of locus and copy number. Here, for the first time, we quantified IME *in vivo*, in terms of protein expression levels, using fluorescent reporter proteins expressed from a single, defined locus in *Caenorhabditis elegans*. To quantify the magnitude of IME, we developed a microfluidic chip-based workflow to mount and image individual animals, including software for operation and image processing. We used this workflow to systematically test the effects of position, number and sequence of introns on two different proteins, mCherry and mEGFP, driven by two different promoters, *vit-2* and *hsp-90*. We found the three canonical synthetic introns commonly used in *C. elegans* transgenes increased mCherry protein concentration by approximately 50%. The naturally-occurring introns found in *hsp-90* also increased mCherry expression level by about 50%. Furthermore, and consistent with prior results examining mRNA levels, protein activity or phenotypic rescue, we found that a single, natural or synthetic, 5’ intron was sufficient for the full IME effect while a 3’ intron was not. IME was also affected by protein coding sequence (50% for mCherry and 80% for mEGFP) but not strongly affected by promoter 46% for *hsp-90* and 54% for the stronger *vit-2*. Our results show that IME of protein expression in *C. elegans* is affected by intron position and contextual coding sequence surrounding the introns, but not greatly by promoter strength. Our combined controlled transgenesis and microfluidic screening approach should facilitate screens for factors affecting IME and other intron-dependent processes.

## INTRODUCTION

Introns are noncoding DNA sequences interspersed between the protein-coding DNA sequences called exons. When a gene is transcribed, the introns are removed by the splicing machinery to generate a spliced messenger RNA (mRNA) (Shi 2017a; Shi 2017b). Introns can affect gene expression in several ways (reviewed in (Jo and Choi 2015)). For example, introns facilitate the creation of different protein isoforms- a phenomenon known as alternative splicing (Roy *et al*. 2013; Carazo *et al*. 2018). Introns can also encode catalytic and regulatory RNAs (Engreitz *et al*. 2016; Wong *et al*. 2016; Bai *et al*. 2017; Hube *et al*. 2017; Joung *et al*. 2017; Steiman-Shimony *et al*. 2018). And, introns may restrict genomic instability (Bonnet *et al*. 2017). Finally, and the focus of this report, introns can increase the expression level of a gene via a number of different mechanisms. The expression level is referred to as intron mediated enhancement of gene expression (IME; reviewed in (Le Hir *et al*. 2003; Gallegos and Rose 2015; Shaul 2017)).

Here, we are focused on IME of gene expression in order to: 1) quantify the magnitude that introns increase protein expression levels in *C. elegans*, and 2) to develop *C. elegans* as an *in vivo*, genetically tractable, screenable model to help reveal the many different ways that introns act to influence the functions of biological systems. This understanding may be particularly important not just for basic biology (Yofe *et al*. 2014) and biotechnology (Lacy-Hulbert *et al*. 2001; Xu *et al*. 2018), but for human diseases, including those where intronic mutations affect gene regulation (Riley and Krieger 2005; Vaz-Drago *et al*. 2017). Thus, employing *C. elegans* to understand the basic biology of introns using *in vivo* measurements of reporter proteins may offer novel insights into the physiology and molecular biology of introns, including IME of gene expression levels, and the ways that allelic variants of introns affect gene expression – both cis and trans.

In general, IME is defined as the increase in expression of an intron-containing gene compared to an otherwise-identical intronless version of the same coding sequence (Brinster *et al*. 1988). There are several mechanisms by which IME is thought to act (Le Hir *et al*. 2003; Gallegos and Rose 2015; Shaul 2017). In mammalian and plant systems, introns have been shown to increase the rate of transcription, and this can be mediated by splicing independent functions of splicing factors such as U1 snRNA (Furger *et al*. 2002; Kwek *et al*. 2002; Damgaard *et al*. 2008) or by gene looping (Moabbi *et al*. 2012). Moreover, proteins deposited onto spliced transcripts called the exon junction complex (EJC) have been shown to increase mRNA transport from the nucleus to the cytoplasm (Le Hir *et al*. 2001; Valencia *et al*. 2008), and might also play a role in increased translation efficiency of spliced transcripts (Wiegand *et al*. 2003; Nott *et al*. 2004; Lee *et al*. 2009).

There are still many unresolved questions about the mechanisms of IME. However, it seems for the most part, and consistent with what we observe in this report, that a 5’proximal intron is sufficient for IME of gene expression level, at least for most intron sequences studied in detail (Nott *et al*. 2003; Rose 2004). One way 5’ proximal introns can increase gene expression levels is by causing more recruitment of RNA polymerase II to the transcription initiation site than would occur with an intronless coding sequence (Das *et al*. 2007; Damgaard *et al*. 2008). Evidence also suggests that promoter-proximal introns affect transcription initiation via chromatin modifications (Parra *et al*. 2011; Bieberstein *et al*. 2012). A 5’ intron is also sufficient to recruit chromatin opening marks (Bieberstein *et al*. 2012). Experiments swapping the sequence and orientation of introns suggest that it is the DNA and not the RNA sequence that is critical for marking chromatin (Rose *et al*. 2011). Rose and Gallegos have put forth a model in which promoter-proximal introns that contain stimulatory signals control transcription initiation or re-initiation, in a chromatin state dependent manner, either by allowing for an open chromatin state or by facilitating activating histone modifications (Rose and Gallegos, 2015).

Previous reports have measured IME of gene expression level in terms of RNA abundance (Rose 2002; Rose 2004; Bieberstein *et al*. 2012), protein activity (Nott *et al*. 2003; Rose 2004) or phenotype (related directly to protein activity (Okkema *et al*. 1993)). These reports found that a 5’ intron was sufficient for IME; however, the best available technology at the time of these reports lacked control of copy number *<u>and</u>* locus, which could have affected the ability of transgenes to be expressed into protein. Additionally, the measurement of protein expression abundance was from biochemical extracts of protein (activity or densitometry-measured abundance of extracted protein), which could have included differences in extraction efficiency. Finally, in some cases, the protein expression was transient (temporally unstable due to the nature of transfection/transduction). Thus, prior reports may have incorporated additional sources of experimental error that we are able to obviate with modern genome editing and *in vivo* quantification of reporter signal.

Most reporter genes created for expression in C. *elegans* include a specific set of introns to achieve IME. The classic vector kit created by Andrew Fire relied on three synthetic introns to boost expression level (Fire 1995). The widespread popularity of this system has had the consequences that the majority of reporter genes in *C. elegans* incorporate these three synthetic introns. Moreover, the software for *in silica* codon optimization of DNA sequences intended for expression in *C. elegans* recommends the usage of these three introns (Redemann *et al*. 2011). However, despite of the widespread adoption and usage of these intron sequences, there is no published data quantifying the effect of these sequences on protein expression level.

Here, we developed a microfluidic/microscopy platform and implemented it to quantify IME in individual live animals. We measured the magnitude of IME *in vivo* using fluorescent reporter proteins expressed from a single defined locus in the genome. Using the C. elegans *hsp-90* promoter to drive expression of an mCherry reporter protein, we found that the classical three synthetic introns, or two natural introns increased protein expression roughly 50%. We varied intron number, sequence, and position in order to determine which introns and positions conferred IME. As has been reported by others, we found that a single 5’-proximal synthetic or natural intron increased gene expression similar to that of 3 synthetic introns, while a more 3’-positioned intron did not. Constructs that contained the stronger *vit-2* promoter showed slightly increased IME. Addition of the classical three synthetic introns to mEGFP reporters increased IME by about 80% compared to intronless mEGFP. Taken together, our results suggest that IME is influenced by intron position and sequence context but is not constrained by free translational capacity. Finally, the microfluidic/microscopy system we developed and implemented here to quantify IME enables researchers to quantify gene expression at a throughput more similar to current worm flow instruments, but, unlike those technologies, it records and image of each animal.

## RESULTS

### A set of *C. elegans* expressing fluorescent proteins with different intron configurations

We designed and constructed a set of eleven different *C. elegans* strains wherein we edited the genomes to place single copy reporter genes at the locus on Chromosome II designated by a transposon insertion, ttTi5605 (Frokjaer-Jensen *et al*. 2008; Frokjaer-Jensen *et al*. 2012; Frokjaer-Jensen 2013; Frokjaer-Jensen *et al*. 2014). We varied the promoter (*hsp-90* or *vit-2*), and/or number, position or sequence of introns (synthetic or natural); we kept the *unc-54* terminator as a constant. The constructs integrated into the genomes of the resulting strains are shown in Figure 1 and listed in Table 1. The sequences of all reporter gene are listed in Supplemental Figure 1. Because autofluorescence levels are significantly lower in mCherry compared with GFP, we used mCherry for the majority of the work, except for when testing whether the protein coding sequence affects IME magnitude.

**Figure 1.**
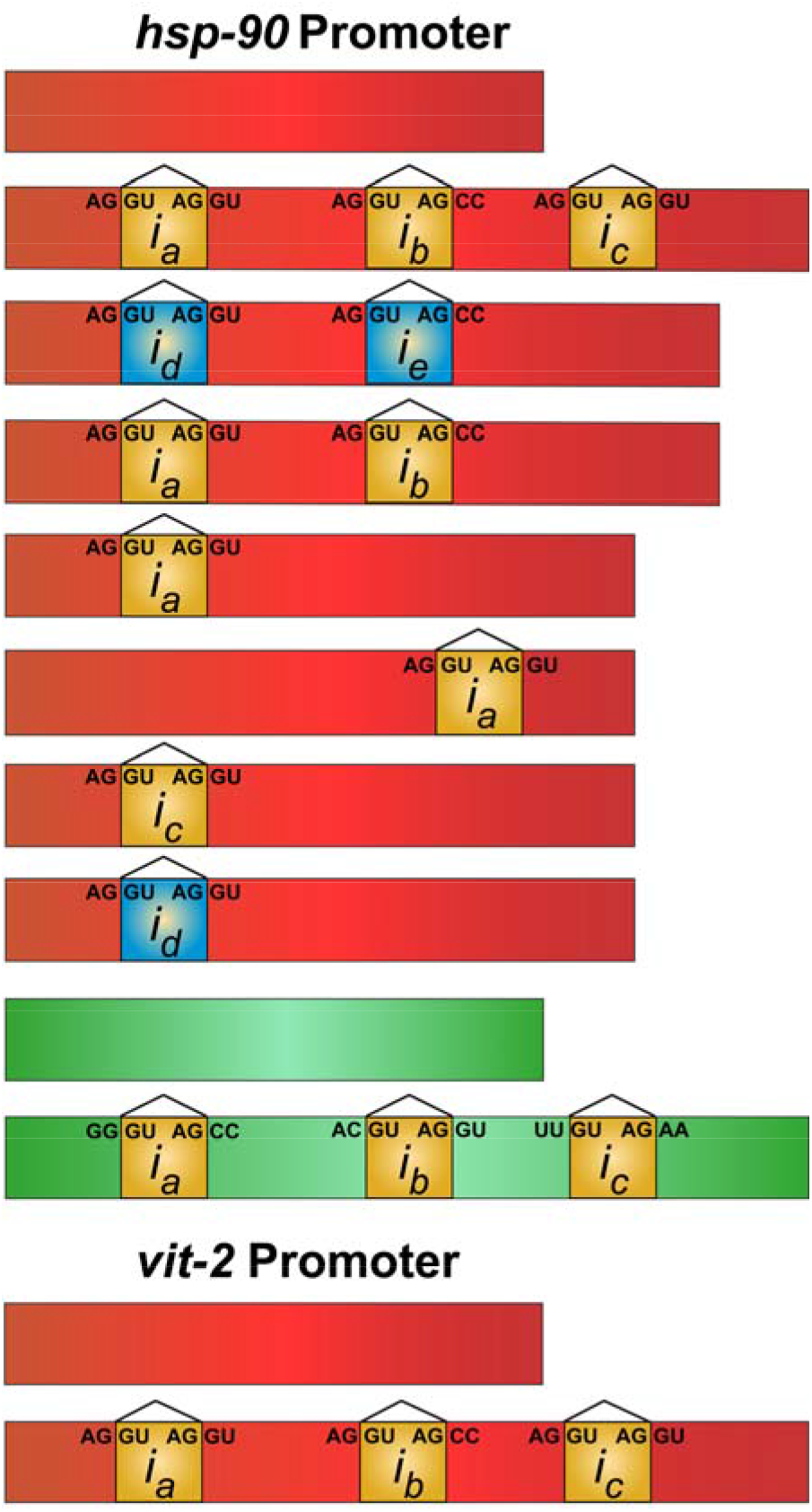
A set of transgenic *C. elegans* expressing fluorescent reporter proteins with and without introns from a single autosomal locus in the genome. Promoter names are listed above the reporter gene coding sequences for which they control the expression, and a graphical representation of each construct is shown on the right. The exons are shown in red, and introns are shown progressing from yellow to green, and labeled *i_a-e_*. Introns *i_a-c_* are the canonical synthetic introns, and *i_d-e_* are the first two introns naturally found in *C. elegans’ hsp-90*. Supplementary Figure 1 provides strain names and Tables 1&2 provide additional information on plasmids and sequences.

**Table 1.**
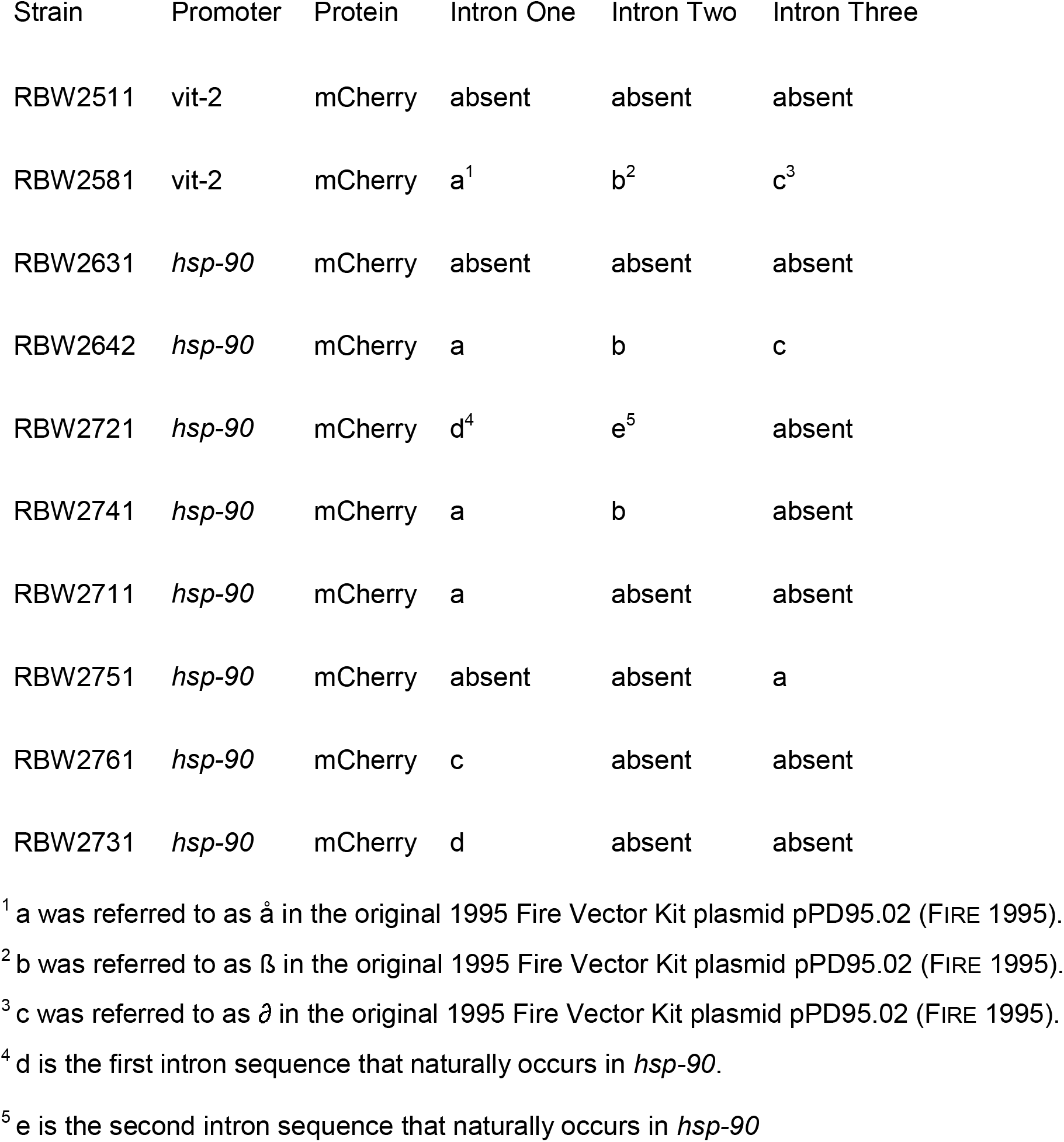

### A cost-effective, semi-automated microfluidic device that quantifies gene expression levels in individual animals

Here we modified a previous microfluidic chip design (Crane *et al*. 2012; San-Miguel *et al*. 2016) to develop an instrument that allowed us to quantify gene expression in whole animals *and* acquire an image of each animal while doing so. In this system, air pressure causes the worms to flow from a pressurized 1.5 mL plastic tube into the PDMS chip, where they are subsequently imaged (Figure 2). On the chip, animals are imaged in a “J” shaped orientation, and then flow out of the imaging chamber into an exit tube, where it can be streamed as waste, or manually captured for further analysis by diverting the exit tube onto solid or liquid worm growth media. Thus, the design can also accommodate manual sorting of animals with complex phenotypes. Figure 2A shows an overview of the setup. Figure 2B details the process of entry and exit into the imaging chamber. Figure 2C shows how we quantify signal from animals. We can typically image about 100 worms in twenty minutes. Measuring only 50 worms takes almost the same amount of time as preparing the device and worms for any amount of animals to measure takes a fixed amount of time, which is variable, depending on experience level. This instrument is not nearly as fast as the Copas Biosort, but, importantly, it captures an image of every worm, and the values are not subject to changes in animal orientation or flow rate.

**Figure 2.**
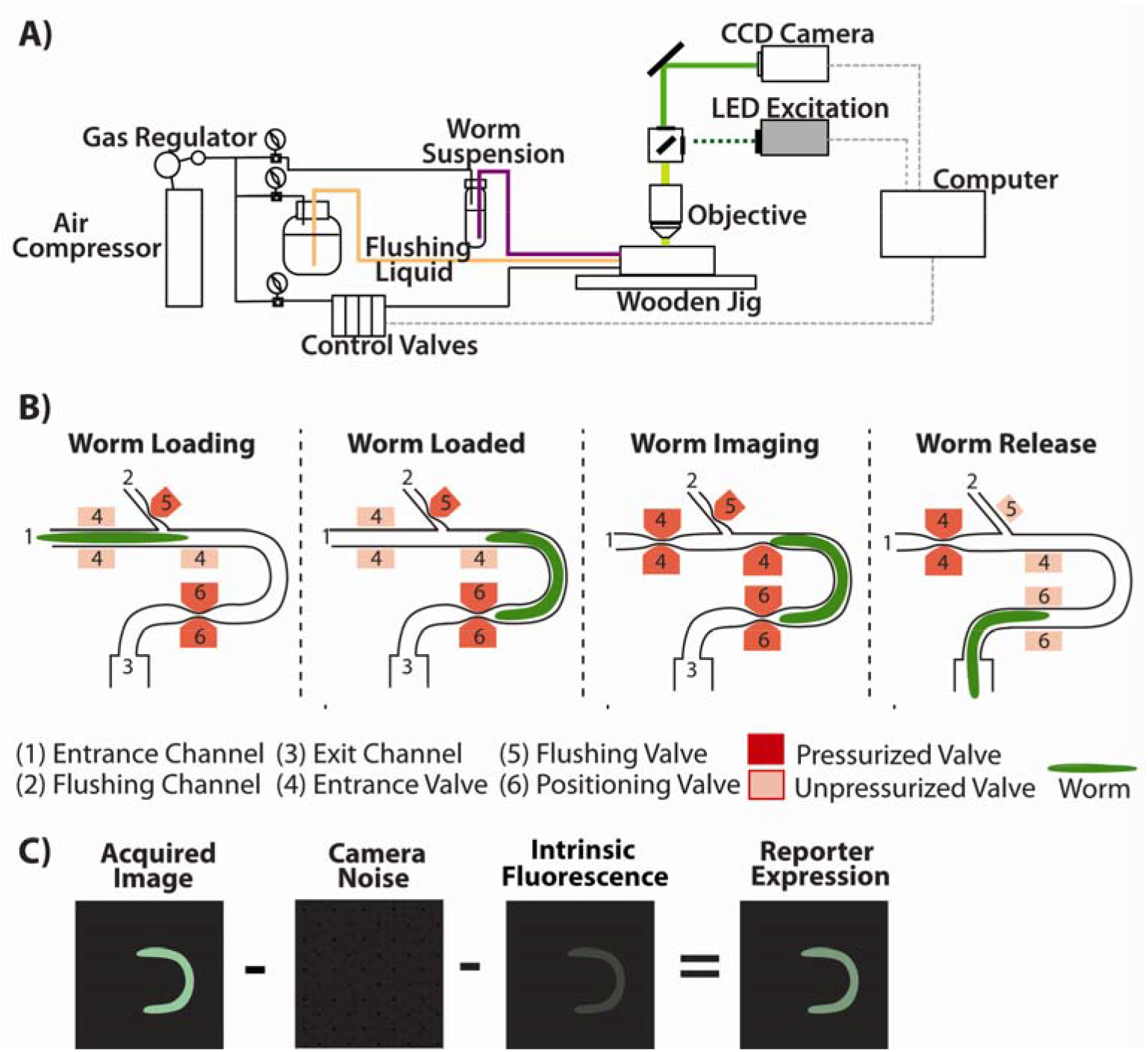
Overview of microfluidic imaging device schematics and image calibration procedure. A) shows a schematic overview of the worm microfluidic measurement device. B) displays a time series of cartoons showing valve openings and closings occurring during imaging of experimental groups of worms. Depth of field on our objective is approximately 55 micrometers (see Materials and Methods), and our imaging chamber is 50 micrometers deeps in z, ensuring we capture all the signal form each animal. C) shows the image correction protocol we used to determine average voxel intensity, quantifying expression level as a function of concentration (not total signal).

### The IME effect of the canonical three synthetic introns on mCherry is an approximately 50% increase in protein expression level

Because of the widespread usage of the three canonical synthetic introns created by Andrew Fire (Fire 1995), we first wanted to determine the IME magnitude using these introns. Here, we used the constitutive, ubiquitously expressed, *hsp-90* promoter to control expression of mCherry with and without these synthetic introns. The *C. elegans* homolog of *HSP90* and is expressed in most/all somatic cells during adulthood. We expressed these reporter genes from the same locus on chromosome II (inserted at ttTi5605; (Frokjaer-Jensen *et al*. 2008; Frokjaer-Jensen *et al*. 2012; Frokjaer-Jensen *et al*. 2014)). By measuring whole animal mCherry signal in our microfluidic chip (after correcting for intrinsic fluorescence in wild-type animals), we found that the presence of the three synthetic introns increases the protein expression level by 46% (Fig 3).

**Figure 3:**
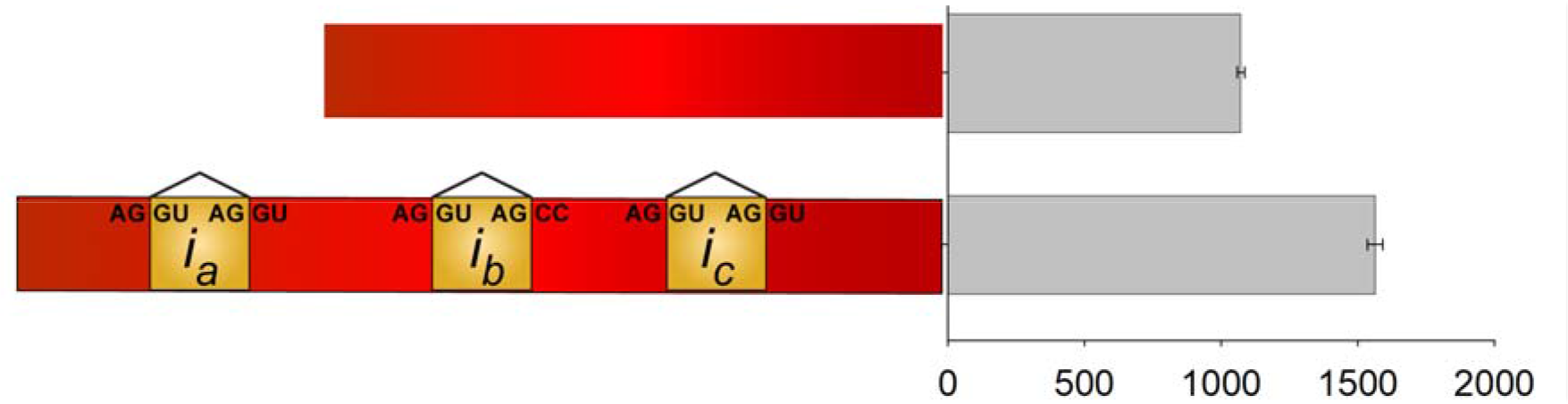
Synthetic introns increase mCherry expression level by about 50%. Figure shows results of experiments measuring mCherry with and without three synthetic intron: under *hsp-90* promoter control in live young adult *C. elegans*. X axis shows average ti counts per voxel. Y axis shows a graphical representation of mCherry with (bottom) and without (top) introns. Error bars show 95% confidence intervals. Three synthetic intron increase expression level by 45.88%; P < 0.001, Mann-Whitney Rank Sum Test. Given tha we measured several different strains compared to the control, we also performed a Kruskal Wallis One Way Analysis of Variance on Ranks followed by Dunn’s Method to compare to the intronless control and detected a significant difference P < 0.05. At least three independen experiments with at least 185 animal were performed (185-297).

### Natural Introns Have Same Effect Size as Synthetic Introns

Because the canonical introns used in *C. elegans* reporter transgenes are synthetic, we wanted to determine whether natural introns affect IME magnitude in similar fashion. To do this, we replaced the synthetic introns in our *P_hsp-90_::mCherry::T_unc-54_* construct with the first two natural *hsp-90* introns. The natural introns were placed at the same exon-intron boundaries as the synthetic introns which maintained the same 5’ and 3’ splice junctions (Table 2). The natural *hsp-90* introns increased protein expression level to the same extent as the three synthetic introns (~50%), shown in Figure 4.

**Figure 4.**
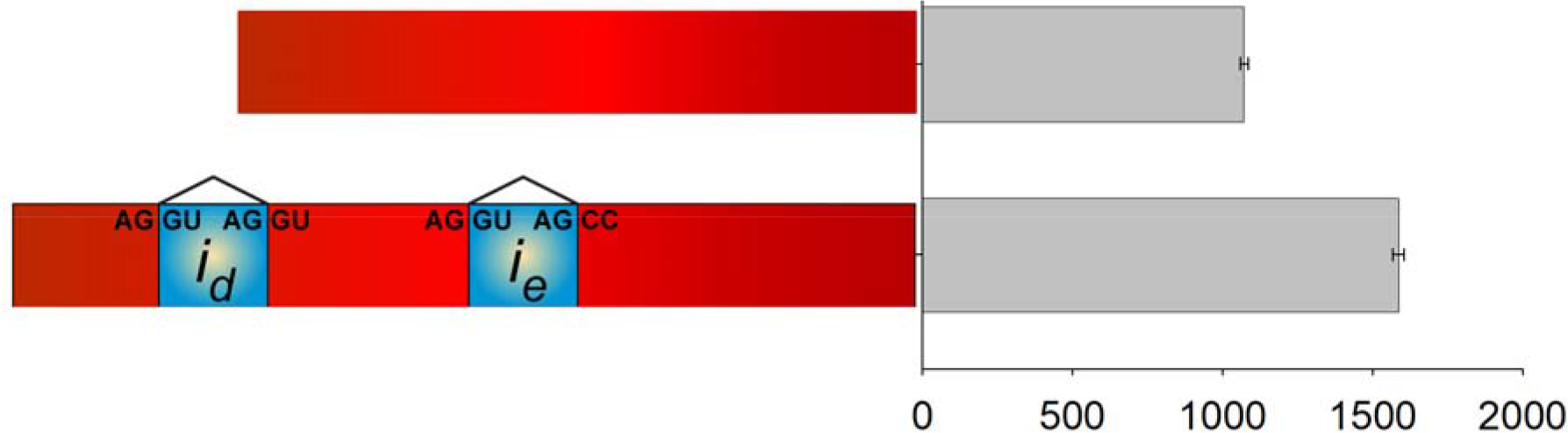
Natural introns provide the same level of IME as synthetic introns. X axis shows average tif counts per voxel. Y axis shows a graphical representation of reporter genes with and without natural introns. Error bars show 95% confidence intervals. Natural introns significantly increase the expression level of mCherry by 47.98%; P < 0.001, Mann-Whitney Rank Sum Test. Given that we measured several different strains compared to the control, we also performed a Kruskal-Wallis One Way Analysis of Variance on Ranks followed by Dunn’s Method to compare to the intronless control and detected a significant difference P < 0.05. At least three experiments with at least 179 total animals were performed (179-297 animals).

### A single 5’ intron produces the same IME effect as three synthetic introns

Prior reports demonstrated that a single 5’ intron was sufficient for some level of IME (Okkema *et al*. 1993; Nott *et al*. 2003; Rose 2004). Thus, we worked to determine how the number and sequence of introns affected IME. To do this, we generated strains containing one, two or three synthetic or natural introns at different positions in the gene, shown in Figure 1. We found that two synthetic introns produced the same IME effect as all three synthetic introns. Strains containing only a single natural or synthetic intron at the first position had the same level of IME as all three introns (Figure 5). A single intron at the 3’ site was not sufficient to increase gene expression.

**Figure 5.**
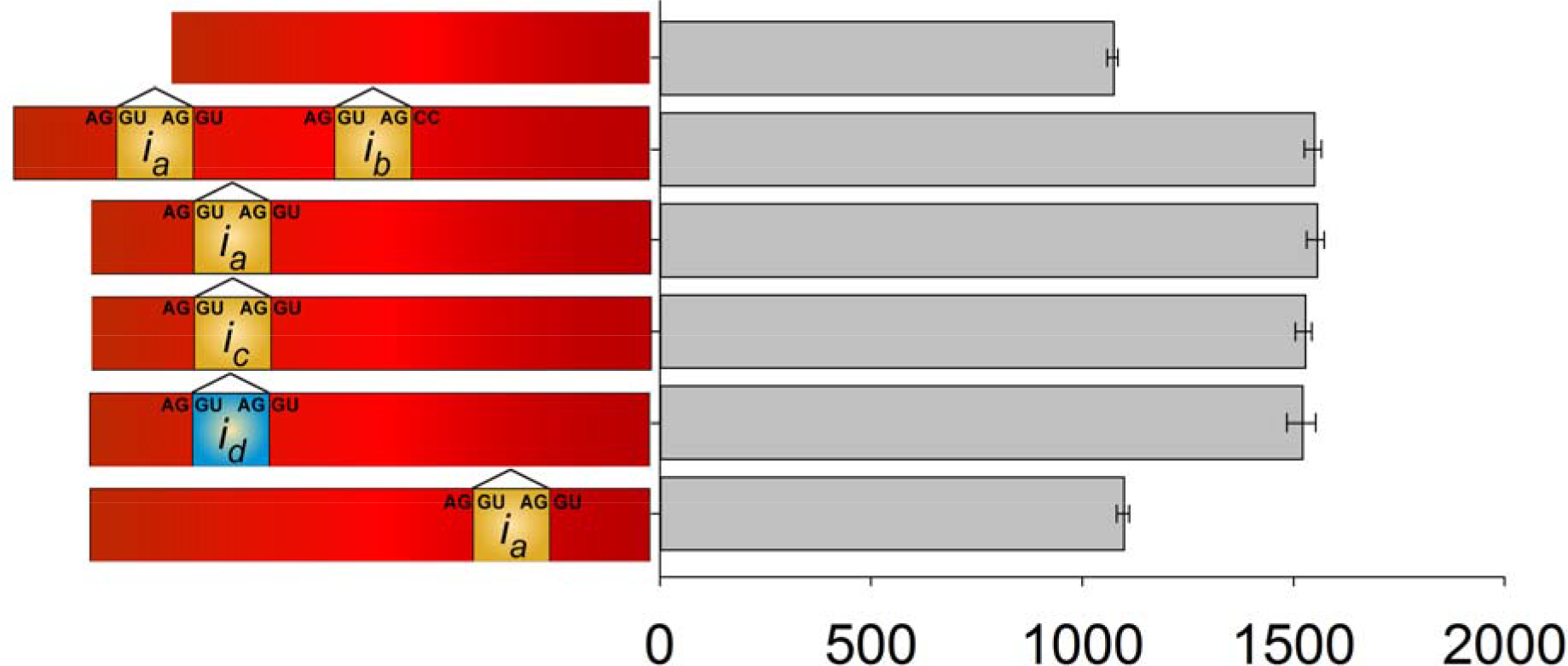
A single natural or synthetic 5’ intron is sufficient for the full magnitude of IME and a single 3’ intron is not. X axis shows average tif counts per voxel. Y axis shows a graphical representation of reporter genes with different intron configurations. Error bars show 95% confidence intervals. With the exception of animals bearing the reporter gene with only *i_a_* at the most 3’ position in the coding sequence, all other intron bearing strains expressed significantly more protein than the intronless strain; Kruskal-Wallis One Way Analysis of Variance on Ranks followed by Dunn’s Method, P < 0.05 for all comparisons. At least three independent experiments with at least 100 total worms per strain (103-298 animals per group).

### IME effect size is only slightly increased under control of one of the strongest promoters in *C. elegans*

To determine if promoter strength affected IME, we replaced the *hsp-90* promoter with *vit-2*, perhaps the strongest promoter in *C. elegans*. The *vit-2* gene encodes a yolk protein expressed in the intestine of *C. elegans*. When considering it is only expressed from the 20 intestine cells (Spieth *et al*. 1988), it is possibly the most highly expressed gene (in terms of concentration) in adulthood (Hillier *et al*. 2009), which is when we measured the animals (day one of adulthood). If the expression boost were primarily constrained by free translational capacity, then the magnitude of protein increase could depend significantly on the transcriptional strength of the promoter. That would mean that stronger promoters would receive less of a boost. We constructed two strains using the *vit-2* promoter to drive mCherry with either no introns, or the three synthetic introns (Figure 1; Table 1). The presence of the three synthetic introns increased protein expression by ~50% (Figure 6) showing that the distinctly regulated, and stronger *vit-2* promoter does not have a large effect on the magnitude of IME at the protein level compared to the *hsp-90* promoter (Figure 6B, 53.55% vs. 45.88%; *P* = 0.001, Kruskal-Wallis One Way Analysis of Variance on Ranks and Dunn’s Method). Note that the absolute value of protein produced under *vit-2* promoter control was much higher. Thus, the magnitude of the IME boost does not appear to be constrained by free translational capacity.

**Figure 6.**
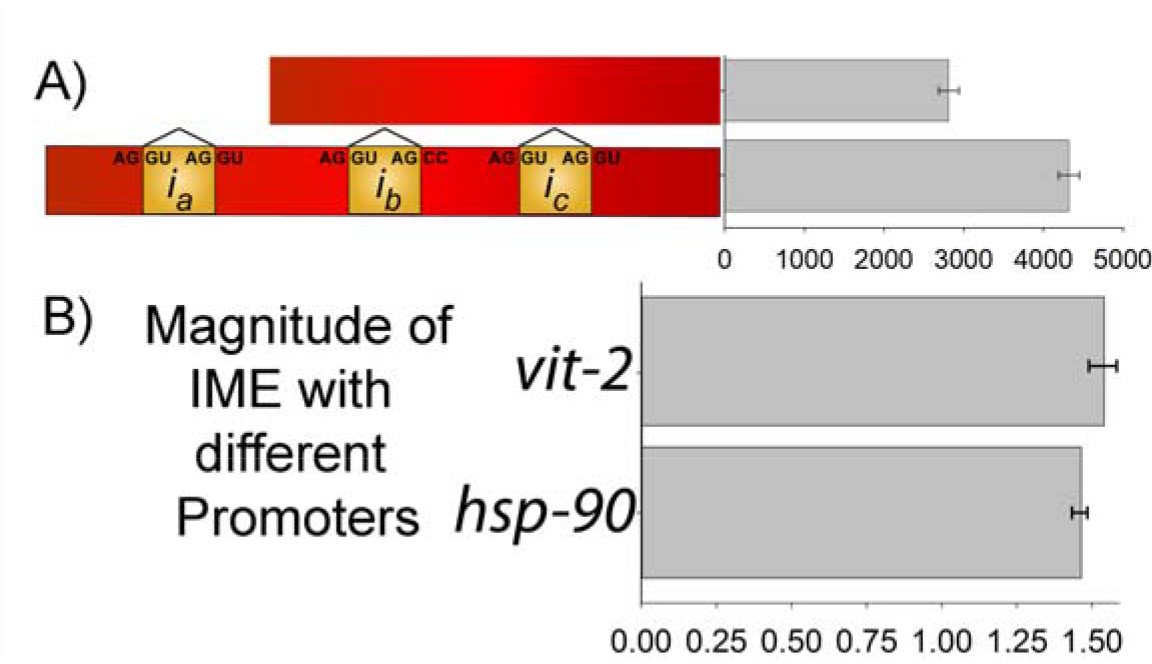
Effects of synthetic introns on expression levels of reporter genes controlled by *vit-2* and *hsp-90* promoters. **A)** shows results of experiments measuring mCherry with and without three synthetic introns under *vit-2* promoter control in live young adult *C. elegans*. X axis shows average tif counts per voxel. Y axis shows a graphical representation of mCherry with (bottom) and without (top) introns. Error bars show 95% confidence intervals. Synthetic introns significantly increase mCherry expression level under *vit-2* control. Mann-Whitney Rank Sum Test, P < 0.001. Results are from three independent experiments with at least 130 total animals per group (134-163). **B)** shows the effect size of IME of mCherry expression level under control of *hsp-90* and *vit-2*. There is a slight but significant increase of IME effect size when mCherry expression is controlled by the much stronger *vit-2* promoter (53.55% vs. 45.88%). P < 0.05, Kruskal-Wallis One Way Analysis of Variance on Ranks and Dunn’s Method. No significant difference was detected between the IME effects size of the natural introns under control of *hsp-90* when compared to the other two strains.

### Protein coding sequence affects the magnitude of IME

We wanted to determine if protein coding sequence would affect the IME effect size for the same three synthetic introns used with mCherry. To do this we generated strains containing the mEGFP coding sequence with and without the introns (see Figure 1). Figure 7 shows that we observed an IME effect size of ~80%, which is significantly different from the approximate 50% increase in gene expression conferred by the same three introns in the context of the mCherry coding sequence.

**Figure 7.**
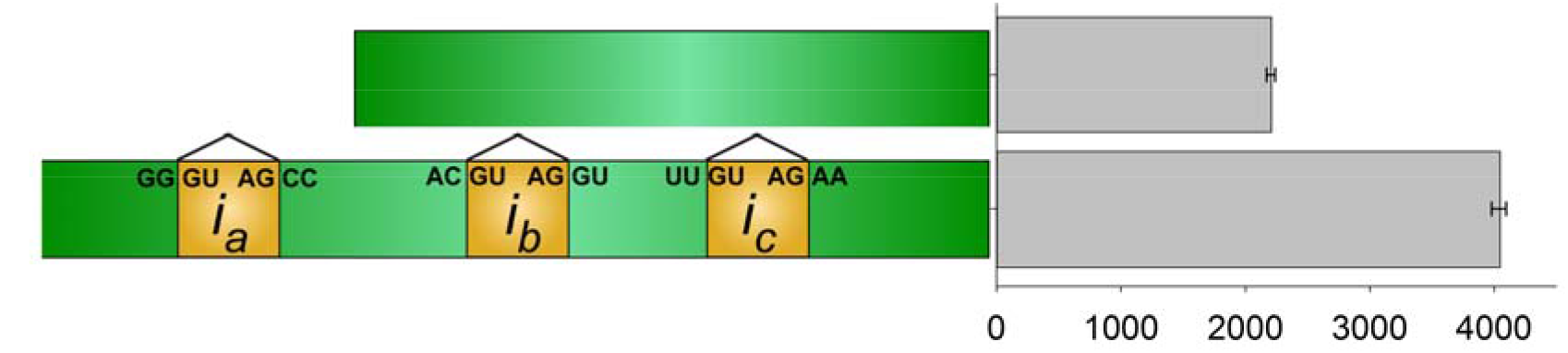
Coding sequence affects the magnitude of IME. X axis shows average tif counts per voxel. Y axis shows a graphical representation of mEGFP-based reporter genes with and without synthetic introns. Error bars show 95% confidence intervals. The same three synthetic introns in mCherry significantly increase mEGFP expression level by 83%. P < 0.001, Two Way ANOVA followed by the Holm-Sidak method for multiple comparison procedures. Results are from three independent experiments measuring at least 200 total animals per group (215-260).

## DISCUSSION

We described *in vivo* means to quantify intron mediated enhancement of gene expression (IME) by fluorescent reporter protein expressed in the tissues of living C. elegans. We found that both promoter strength and contextual coding sequence can affect the level of IME. A single 5’ intron gave the same IME as a “classical” set of three synthetic introns used for *C. elegans* gene expression, and the same IME as the first two natural introns of *hsp-90*. Our results demonstrate that splicing of a 5’ proximal intron is necessary and sufficient for IME. To our knowledge, these experiments comprise the first quantitative study of IME in a living animal.

Our findings corroborate the results from prior studies showing that a single 5’ intron was sufficient for IME of gene expression using phenotype (Okkema *et al*. 1993), extracted RNA (Rose 2004) or extracted protein activity (Nott *et al*. 2003; Rose 2004) as readouts for the effect of introns on gene expression level. Our results are consistent with the idea that IME of protein expression occurs via transcription, translation, or both. They are consistent with a transcriptional mechanism in which the splicing machinery at the 5’ intron rerecruits RNA polymerase back to the transcriptional start site. They are consistent with the possibility that a 5’ intron may affect transcription by opening chromatin, for instance, by recruitment of the molecular machinery that makes access-permissive histone modifications to promoter-proximal histones. Given cotranscriptional splicing (Kotovic *et al*. 2003; Listerman *et al*. 2006), our results are also consistent with the possibility that a 5’ intron can affect translation via proteins deposited during splicing onto the intron-exon junction (called the exon junction complex or EJC) which can recruit the translation machinery to EJC-marked transcripts (Cheng *et al*. 2006).

### Using *C. elegans* to study intron function

Much of the prior work on IME has relied on quantification of mRNA and enzymatic activity in lysates of *Arabidopsis* and of vertebrate cells cultured *in vitro*. Since introns in *C. elegans* are similar to introns in humans in terms of size (mode is 47nt for *C. elegans*, 87 for *H. sapiens*, and even at least one very large intron > greater than 100,000nt as might be found in humans (Spieth and Lawson 2006)), we are optimistic that our ability to quantify IME *in vivo* in living *C. elegans* will allow studies of intron biology relevant to understanding IME in vertebrates.

Our ability to study IME in cells of living animals will allow use of previously developed (Mendenhall *et al*. 2015) highly reproducible confocal microscopic techniques to quantify intron function in particular cells and tissues. The higher throughput methods described here should allow investigators to take even better advantage of some of the strengths of this model system; its optical transparency, its short life cycle, its full editable genome, and its eminent utility in forward and reverse genetic screens. Moreover, the recent development of synthetic food medium as an alternative to the typically used OP50 *E. coli* strain has should allow execution of chemical genetic screens, using for example, FDA approved pharmaceuticals packaged into liposomes for delivery into the worm body via the alimentary canal (Flavel *et al*. 2018).

### Quantifying IME by imaging reporter proteins *in vivo*

The methods described here allow us to quantify whole animal reporter gene signals from live *C. elegans* at medium throughput (30-100 animals in 30 minutes). Our previous measurements of gene expression in whole animals (Mendenhall *et al*. 2012; Mendenhall *et al*. 2015; Mendenhall *et al*. 2017a; Mendenhall *et al*. 2017b)) utilized a higher throughput (300-1000 animals in similar timeframe) flow sorting device (COPAS Biosorter, Union Biometrica) that measures signal from animals as they pass by a photomultiplier tube. This instrument does not acquire images of each animal, forgoing acquisition of potentially useful image features. By contrast, our system collects an image of each physically restricted animal, oriented in the same J shaped orientation. The system’s objective lens has a large depth of field, (> 50 μm) allowing collection of signal from the entire depth in z of the immobilized worm.

### How Much Can IME Increase Expression Level?

Prior careful, methodological studies of IME of gene expression levels have revealed much about the mechanisms by which introns act to increase expression levels. However, more work still needs to be done in order to answer many lingering questions about IME (Le Hir *et al*. 2003; Gallegos and Rose 2015; Heyn *et al*. 2015; Shaul 2017). Frankly, it can be technically difficult to separate the contributions of individual genes to affect the seemingly coordinated and possibly overlapping intracellular events occurring to cause IME. The events occurring to produce IME seem likely to utilize proteins that are essential for development and homeostasis – which, minimally, include transcription, splicing, nuclear export and translation machinery. Nevertheless, this and prior studies have quantified the effect size of IME of gene expression level through a variety of means. Below, we discuss the range of increases in gene expression that introns can cause. We are aware of much early work with SV40 introns, but here we focus our discussion on more recent, 21^st^ century reports that more precisely quantified the effects of introns on gene expression levels.

In all the studies of IME, reporter genes with no introns were compared to other reporter genes that bore one or more introns. We found that introns increased protein expression level by about 50% or 100%, depending on coding sequence, and that a single 5’ intron was sufficient for the full IME effect. In Nott et al., 2003, using transient transfections of HeLa cells, they found that a single 5’ intron was sufficient to increase gene expression, measured by *Renilla* luciferase activity, by 29-fold. In Bieberstein et al. 2012, using BAC recombineering of HeLa cells, they found that the mouse FOS gene expressed 5 fold more RNA (500% more) when it included three natural mouse introns; they also found that a single 5’ intron was sufficient to cause the chromatin markings associated with increased gene expression level. In Rose 2002, he found that several distinct *Arabidopsis* introns increased the expression of reporter gene RNA between 40% and 1490% using single copy reporters integrated randomly throughout the genome of *Arabidopsis*. In Rose 2004, he found that another variety of single 5’-proximal *Arabidopsis* introns were sufficient for increased gene expression level, quantified by both mRNA and extracted protein activity (GUS activity), ranging from 10% to 2000%. Thus, the effects we measured fall between the effects that have been measured before. There are still a lot more introns to test in *C. elegans*. It is likely that future studies will identify *C. elegans* introns capable of increasing gene expression levels even more.

### Understanding the role of introns in human disease

Many previous reports have shown that introns influence human diseases. These diseases fall into two broad classes: heritable diseases (Riley and Krieger 2005; Swarbrick *et al*. 2011; Vaz-Drago *et al*. 2017) and cancers (Hsu *et al*. 2015; Jung *et al*. 2015; Wiesner *et al*. 2015; Neri *et al*. 2017; Lee *et al*. 2018). The intron-dependent processes that are affected often unknown, poorly understood, or if understood, lack experimental means to determine how to suppress the intron-related pathology. For example, for a bipolar risk allele, STRs in the third intron of the calcium encoding CACNA1C gene seem to affect IME of gene expression level (Riley and Krieger 2005). However, the mechanism by which this relatively large, relatively 3’ intron acts to increase gene expression level remain unknown. Thus it is our hope that this screenable *in vivo* system will be able to provide insight into intron-related disease mechanisms, and eventually lead to therapeutic treatments to alleviate human diseases.

## Supporting information

## Author contributions

AM, MC and BS concocted the experimental approach. AM and BS designed and constructed strains in the RB lab. MC and AM designed and constructed the microfluidic device. SY, MC, CB and BJ performed the experiments. MC, BS and AM analyzed the data. AM, RB and MK provided funding. All authors were involved in manuscript writing.

## Funding

Funding was provided by National Institute on Aging grants R00AG045341 to A.M, P50 AG005136 to MK, and a Pilot Grant from the Nathan Shock Center for Excellence in the Basic Biology of Aging at the University of Washington to AM (NIA Grant P30AG013280 to MK). Training grant T32AG000057 supported MC. Work by AM and BS was also funded by National Institute of General Medical Science grant R01GM97479 to RB. This work was also funded by the National Cancer Institute grant R01CA219460 to A.M.

## Acknowledgements

Some strains were provided by the CGC, which is funded by NIH Office of Research Infrastructure Programs (P40 OD010440).

## Materials and Methods

### Strains and Culture Conditions

Here, we used wild-type N2 strain of *C. elegans*, RBW6699 and the strains we listed in Figure 1 and Table 1: RBW2642, RBW2661, RBW2511, RBW2581, RBW2631, RBW2651, RBW2721, RBW2741, RBW2731, RBW2761, RBW2711, and RBW2751; each strain has an insert number corresponding to its strain number, meaning RBW2511 contains hutSi2511. Additional strain details are shown in Supplemental Figure 1. We maintained all strains in 10cm petri dishes on NGM seeded with OP50 *E. coli* in an incubator at 20°. Strains were maintained for at least three generations at 20° in a well-fed state before experimentation. For experiments, animals were synchronized via alkaline hypochlorite treatment followed by overnight hatchout in sterile S-basal to cause entry into the L1 diapause, described in detail in (Mendenhall *et al*. 2015). Synchronous cohorts of animals were grown to gravid adulthood (72 hours at 20° on NGM after the L1 diapause) at a density of approximately 175 animals per 10cm NGM plate seeded with 1mL of OP50 *E. coli* from an overnight 500mL or 1000 mL LB growth culture.

### Generation of transgenic nematodes

Here to make the DNA used for the transgenes we inserted, we used a yeast homologous recombination based cloning system we described recently to generate DNA repair templates (Sands *et al*. 2018). To make the reporter genes, we used 2kb upstream of the ATG for the *hsp-90* promoter and 4kb upstream of the ATG for the *vit-2* promoter. We used the same *unc-54* terminator in all strains (Sands *et al*. 2018). For the pieces of DNA in which we varied introns, we created those with overlap extension PCR and then used those fragments in our yeast HR cloning system. These DNA repair templates were then used in microinjection-based MosSCI transgenesis to insert these reporter genes into the genome at the ttTi5605 Mos transposon locus on Chromosome II in strain RBW6699, which is an outcrossed version of EG6699 containing *unc-119(ed3)*. We injected constructs @ 50ng/μL of repair template + coinjection and selection markers as described previously in (Frokjaer-Jensen *et al*. 2008; Frokjaer-Jensen *et al*. 2012; Frokjaer-Jensen *et al*. 2014)). We sequenced the reporter sequences of all DNA constructs to ensure their sequence integrity and genotyped animals to verify insertion at the ttTi5605 locus. Thus, in this way, we generated all the strains listed in Figure 1 and Table 1, with the exception of RBW2642 and RBW2661, which were previously reported in (Mendenhall *et al*. 2015).

### Microfluidic Measurements of Whole-worm Gene Expression Levels

To prepare animals for entry into the microfluidic imaging device, we washed them from their growth plates into 1.5 mL Eppendorf tubes with filter-sterilized S-basal (to remove particulate matter) containing 0.01% tween (to prevent worms from sticking to plastic). We allowed animals to settle for 5-10 minutes in a 50mL conical vial, and then removed the supernatant, and added fresh S-basal. We repeated this process three times to ensure that that any dust or embryos washed from the plates were removed prior to use. We then placed a “slurper” tube into the 1.5 mL Eppendorf tube. After we sealed and pressurized the Eppendorf tube and slurper lid, animals flowed into the slurper tube, and we manually loaded animals, one at a time, into the imaging chamber of the microfluidic device. We measured all strains, one worm at a time. To quantify signal from the animals in microfluidic device, we mounted the device on a wooden jig under a LED-illuminated, fluorescent Leica MFC165 stereoscope (Leica Microsystems, Wetzlar, Germany) set to 12X magnification, equipped with a 1.0x Planapo objective and a Lumenera 3-3UR camera with a 0.63X optical coupler, in a room at approximately 20°. With this setup our depth of field was approximately 55 micrometers and our xy resolution was approximately 2.7 micrometers. We focused the image on the approximate central plane of the worm body by focusing on the grinder and lumen of the pharynx, shown in Figure 1 in (Russell *et al*. 2017). Worms were physically constrained by the 65×50 micrometer dimensions of the imaging chamber preventing them from moving their bodies out of plane. We measured between 30 and 100 animals per strain per experiment. We manually controlled the flow into the imaging and image acquisition in real time on the computer using Matlab and Micromanager, respectively.

### Microfluidic worm measurement device construction

Devices are constructed as previously detailed in (Crane *et al*. 2012; San-Miguel *et al*. 2016), using standard soft-lithography practices. Briefly, a microfluidic master mold was constructed in the University of Washington Nanofabrication facility using a transparency mask from OutputCity. The AutoCAD^®^ design files are available at GitHub at https://github.com/nomadcrane/elegansImaging. The mold was made using SU8 2025 and following fabrication was coated with a monolayer of silane to prevent PDMS adhesion. The individual microfluidic devices were constructed using a two layer fabrication process similar to (San-Miguel *et al*. 2016) wherein a very small amount high pre-polymer to cross-linker (20:1) ratio of Sylgard 184 was poured on the microfluidic mold sufficient to cover the mold with 1-4mm of polymer. This 20:1 ratio has a very low Young’s Modulus which allows the side valve mechanism to work. This base layer is partially cured for 45 minutes in a 70 °C oven, and then a volume of 10:1 (pre-polymer:cross-linker) Sylgard 184 is poured on top. Because the low Young’s Modulus layer is prone to damage or tearing due to the limited polymer cross-linking, this second layer provides much needed structural support for inlet hole punching and bonding. This stability layer is poured to a thickness of 15-20 mm.

Following curing of the stability layer (2+ hours in 70 °C oven), devices are cut and removed from the mold. Holes for media and valves were punched using a 1mm diameter biopsy punch. To ensure that PDMS debris was not introduced to the flow channels as a result of the hole punching, the quality of the biopsy punch was inspected prior to use, and holes were only punched cleanly once. All holes were punched from the device side through the low Young’s Modulus flow layer. Devices were cleaned using scotch tape prior to bonding to slide glass. Bonding was performed using a Harrick PDC-001 plasma cleaner. Devices and slide glass were exposed to high energy plasma for 35 seconds, and then the device was placed on the slide glass. Bonded devices were allowed to rest for at least 1h in the 70 °C oven prior to use.

To prevent diffusion of gas from the pressurized valves into the flow channels, valves were filled with water prior to use. This was done by connecting tubes containing DI water to the valve channels, and pressurizing the channels. The flushing media for the device used to push out the worms uses filtered M9 media containing 0.01% Triton X-100. To ensure that there was no cross-contamination between samples, the flushing media was used at moderate pressure for 5 min to rinse the device and the loading tubes. Loading and flushing pressures were optimized for each experimental run, but were typically within the range of 1-9 psi. The valve pressures were similarly optimized for individual variation in devices, but typically were used in the range of 45-60 psi.

### Valve Controller Box Fabrication

To control both the valve pressures, and whether the valves are pressurized or released, we developed a control box composed of pressure regulators, solenoid valves, and a USB switch to provide computerized control of all on-chip valves. This system was modeled on earlier systems (Chung *et al*. 2008; Crane *et al*. 2012; San-Miguel *et al*. 2016). A complete list of parts is available on GitHub at https://github.com/nomadcrane/elegansImaging.

### Valve Control Software

Computerized control of both the microfluidic valve control box and the microscope camera was implemented using custom written Matlab^®^ software. To provide generalizability, all interaction with microscope and camera was implemented using micromanager, and controlled through Matlab®. The code, is freely available on GitHub at https://github.com/nomadcrane/elegansImaging.

### Quantification of Whole Animal Fluorescence

Image segmentation was automated to ensure consistency in quantification of expression level between different strains. Images from all strains for the same promoter, fluorescent protein combination were loaded into memory, and an appropriate segmentation threshold that minimized intraclass variance was selected using Otsu’s threshold (Otsu 1979). Following subtraction of the camera noise from image, all animals containing transgenes with the same promoter/fluorescent protein combination were segmented using the same threshold. Segmented images were then opened to remove any spurious pixels. This whole animal segmentation provided consistent size measurements between strains, and had a high degree of accuracy when compared with manual segmentation of the brightfield images.

### Microscope corrections and calibrations

There are multiple sources of noise and variation that are generated during conventional epifluorescent imaging. To correct for individual pixel variation in dark current noise in the CCD camera, we acquired 1,000 images using the same exposure times and binning, but without light directed to the camera. For each pixel, the median value was determined and used as the dark current pixel noise, and was subtracted from each image for all experiments. To correct for intrinsic autofluorescence, we imaged approximately 100 wild-type N2 animals using the two image acquisition settings (excitation/emission filters and camera exposure time) we used to image animals expressing mCherry or mEGFP. The whole animal auto-fluorescence was determined using both the mEGFP and mCherry parameters. There is a distribution of autofluorescence levels within the wild-type population, but for simplicity the population mean autofluorescence levels for both the mEGFP and mCherry fluorescent excitation/emission channels were calculated. These mean values were then subtracted from the transgene expressing strains to determine the magnitude of fluorescence from the fluorescent protein.

For those who care to undertake these endeavors comparing mCherry and mEGFP, we note that the intrinsic fluorescence signals emanating from secondary lysosomes and other intrinsically fluorescent (autofluorescent) bio molecules in wild-type N2 animals (Clokey and Jacobson 1986; Mendenhall *et al*. 2015) are relatively stronger when excited and imaged with mEGFP optics (and thus a greater part of the total signal measured under those optics), compared to the same images acquired using an mCherry filter set. Because of this increased background effect, to accurately determine the level of fluorescence derived from the GFP transgene, it is critical to calculate and then remove the level of autofluorescence from the imaged cells (Lichten *et al*. 2014) (or circumvent it by measuring an area with relatively pure reporter signal (Mendenhall *et al*. 2015)).

---

**Supplementary Figure 1.**
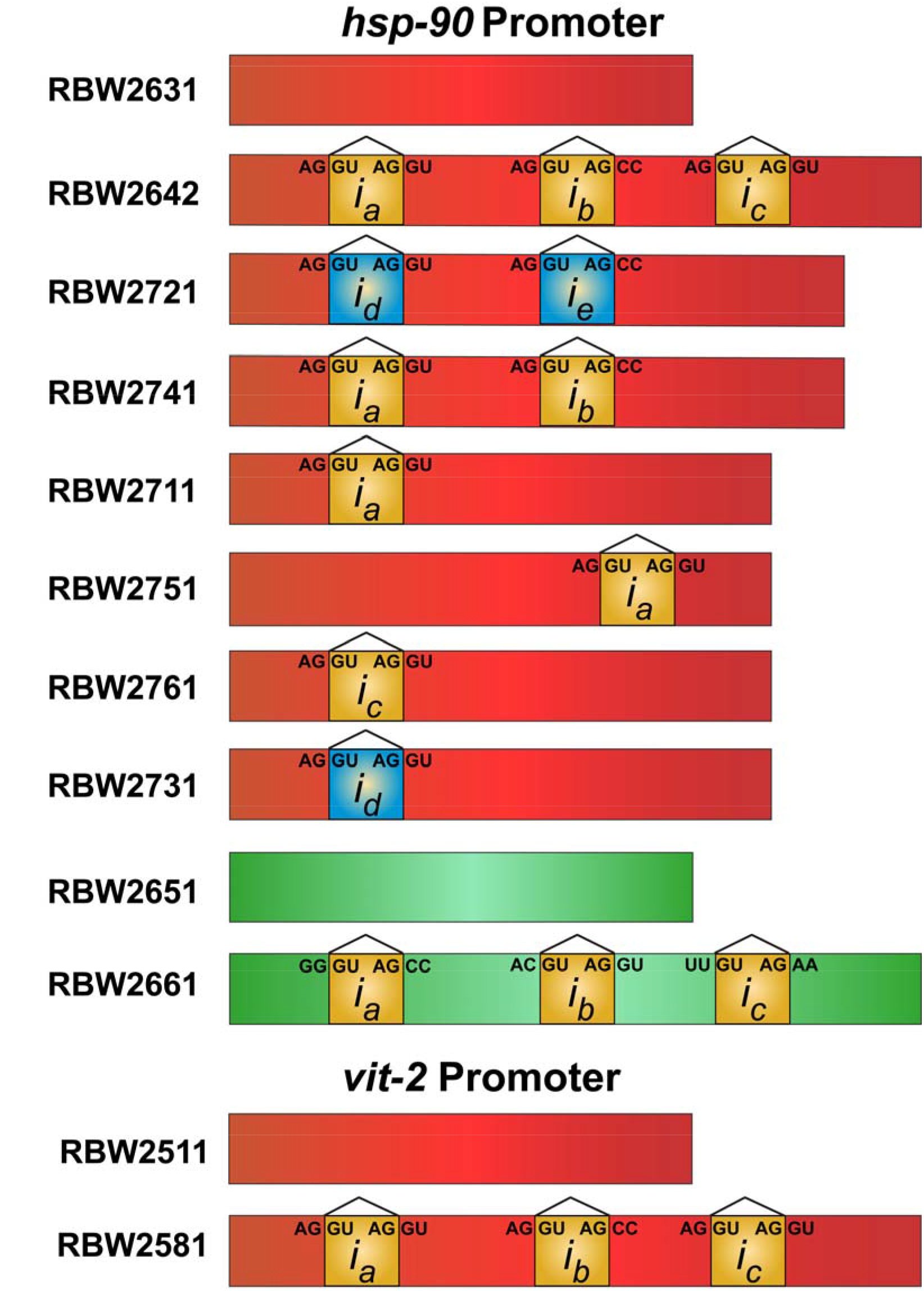
Graphical representation of *C. elegans* strains with and without introns, showing strain name to the left of the graphical representation of the coding sequence of the transgene it bears. Promoters controlling each transgene are listed above the transgenes they control. All strains express the transgenes from the ttTi5605 locus on Chromosome II. All strains use the *unc-54* terminator sequence

