## Supplementary material for "*In vivo* measurements reveal a single 5’ intron is sufficient to increase protein expression level in *C. elegans*"

Supplemental Figure X

DNA sequences of reporter genes.

Coding sequence in Caps. Introns in lower case. Stop codons omitted.

mCherry no introns

ATGGTCTCAAAGGGTGAAGAAGATAACATGGCAATTATTAAAGAGTTTATGCGTTTCAAGGTGCATATGGAGGGATCTGTCAATGGGCATGAGTTTGAAATTGAAGGTGAAGGAGAAGGCCGACCATATGAGGGAACACAAACCGCAAAACTAAAGGTAACTAAAGGCGGACCATTACCATTCGCCTGGGACATCCTCTCTCCACAGTTCATGTATGGAAGTAAAGCTTATGTTAAACATCCGGCAGATATACCAGATTATTTGAAACTTTCATTCCCGGAGGGTTTTAAGTGGGAACGCGTAATGAATTTTGAAGACGGAGGAGTTGTTACAGTGACGCAAGACTCAAGCCTCCAAGATGGAGAATTTATTTATAAAGTCAAACTTCGAGGAACGAATTTCCCCTCGGATGGACCTGTTATGCAGAAGAAGACTATGGGATGGGAAGCTTCAAGTGAAAGAATGTACCCTGAAGACGGTGCTCTTAAGGGAGAGATTAAACAACGTCTTAAATTGAAAGATGGAGGACATTACGATGCTGAGGTGAAGACAACTTACAAAGCCAAAAAACCAGTTCAGCTGCCAGGAGCGTACAATGTTAATATTAAACTGGATATCACCTCCCACAACGAGGATTACACTATCGTTGAGCAATATGAAAGAGCTGAAGGGCGGCACTCGACAGGTGGCATGGATGAATTGTATAAG

mCherry 3 synthetic introns (ia, ib, ic)

ATGGTCTCAAAGGGTGAAGAAGATAACATGGCAATTATTAAAGAGTTTATGCGTTTCAAGGTGCATATGGAGGGATCTGTCAATGGGCATGAGTTTGAAATTGAAGGTGAAGGAGAAGGCCGACCATATGAGGGAACACAAACCGCAAAACTAAAGgtaagtttaaacatatatatactaactaaccctgattatttaaattttcagGTAACTAAAGGCGGACCATTACCATTCGCCTGGGACATCCTCTCTCCACAGTTCATGTATGGAAGTAAAGCTTATGTTAAACATCCGGCAGATATACCAGATTATTTGAAACTTTCATTCCCGGAGGGTTTTAAGTGGGAACGCGTAATGAATTTTGAAGACGGAGGAGTTGTTACAGTGACGCAAGACTCAAGgtaagtttaaacagttcggtactaactaaccatacatatttaaattttcagCCTCCAAGATGGAGAATTTATTTATAAAGTCAAACTTCGAGGAACGAATTTCCCCTCGGATGGACCTGTTATGCAGAAGAAGACTATGGGATGGGAAGCTTCAAGTGAAAGAATGTACCCTGAAGACGGTGCTCTTAAGGGAGAGATTAAACAACGTCTTAAATTGAAAGATGGAGGACATTACGATGCTGAGgtaagtttaaacatgattttactaactaactaatctgatttaaattttcagGTGAAGACAACTTACAAAGCCAAAAAACCAGTTCAGCTGCCAGGAGCGTACAATGTTAATATTAAACTGGATATCACCTCCCACAACGAGGATTACACTATCGTTGAGCAATATGAAAGAGCTGAAGGGCGGCACTCGACAGGTGGCATGGATGAATTGTATAAG

mCherry 2 hsp90 introns (id, ie)

ATGGTCTCAAAGGGTGAAGAAGATAACATGGCAATTATTAAAGAGTTTATGCGTTTCAAGGTGCATATGGAGGGATCTGTCAATGGGCATGAGTTTGAAATTGAAGGTGAAGGAGAAGGCCGACCATATGAGGGAACACAAACCGCAAAACTAAAGgtttgttttttcgcttctgagtcaattttttaaaatatcggttttagGTAACTAAAGGCGGACCATTACCATTCGCCTGGGACATCCTCTCTCCACAGTTCATGTATGGAAGTAAAGCTTATGTTAAACATCCGGCAGATATACCAGATTATTTGAAACTTTCATTCCCGGAGGGTTTTAAGTGGGAACGCGTAATGAATTTTGAAGACGGAGGAGTTGTTACAGTGACGCAAGACTCAAGgtatttttagttttaattattttatggcaataattccattaatttcagCCTCCAAGATGGAGAATTTATTTATAAAGTCAAACTTCGAGGAACGAATTTCCCCTCGGATGGACCTGTTATGCAGAAGAAGACTATGGGATGGGAAGCTTCAAGTGAAAGAATGTACCCTGAAGACGGTGCTCTTAAGGGAGAGATTAAACAACGTCTTAAATTGAAAGATGGAGGACATTACGATGCTGAGGTGAAGACAACTTACAAAGCCAAAAAACCAGTTCAGCTGCCAGGAGCGTACAATGTTAATATTAAACTGGATATCACCTCCCACAACGAGGATTACACTATCGTTGAGCAATATGAAAGAGCTGAAGGGCGGCACTCGACAGGTGGCATGGATGAATTGTATAAG

mCherry 2 synthetic introns (ia, ib)

ATGGTCTCAAAGGGTGAAGAAGATAACATGGCAATTATTAAAGAGTTTATGCGTTTCAAGGTGCATATGGAGGGATCTGTCAATGGGCATGAGTTTGAAATTGAAGGTGAAGGAGAAGGCCGACCATATGAGGGAACACAAACCGCAAAACTAAAGgtaagtttaaacatatatatactaactaaccctgattatttaaattttcagGTAACTAAAGGCGGACCATTACCATTCGCCTGGGACATCCTCTCTCCACAGTTCATGTATGGAAGTAAAGCTTATGTTAAACATCCGGCAGATATACCAGATTATTTGAAACTTTCATTCCCGGAGGGTTTTAAGTGGGAACGCGTAATGAATTTTGAAGACGGAGGAGTTGTTACAGTGACGCAAGACTCAAGgtaagtttaaacagttcggtactaactaaccatacatatttaaattttcagCCTCCAAGATGGAGAATTTATTTATAAAGTCAAACTTCGAGGAACGAATTTCCCCTCGGATGGACCTGTTATGCAGAAGAAGACTATGGGATGGGAAGCTTCAAGTGAAAGAATGTACCCTGAAGACGGTGCTCTTAAGGGAGAGATTAAACAACGTCTTAAATTGAAAGATGGAGGACATTACGATGCTGAGGTGAAGACAACTTACAAAGCCAAAAAACCAGTTCAGCTGCCAGGAGCGTACAATGTTAATATTAAACTGGATATCACCTCCCACAACGAGGATTACACTATCGTTGAGCAATATGAAAGAGCTGAAGGGCGGCACTCGACAGGTGGCATGGATGAATTGTATAAG

mCherry 1 synthetic intron 5’ end (ia)

ATGGTCTCAAAGGGTGAAGAAGATAACATGGCAATTATTAAAGAGTTTATGCGTTTCAAGGTGCATATGGAGGGATCTGTCAATGGGCATGAGTTTGAAATTGAAGGTGAAGGAGAAGGCCGACCATATGAGGGAACACAAACCGCAAAACTAAAGgtaagtttaaacatatatatactaactaaccctgattatttaaattttcagGTAACTAAAGGCGGACCATTACCATTCGCCTGGGACATCCTCTCTCCACAGTTCATGTATGGAAGTAAAGCTTATGTTAAACATCCGGCAGATATACCAGATTATTTGAAACTTTCATTCCCGGAGGGTTTTAAGTGGGAACGCGTAATGAATTTTGAAGACGGAGGAGTTGTTACAGTGACGCAAGACTCAAGCCTCCAAGATGGAGAATTTATTTATAAAGTCAAACTTCGAGGAACGAATTTCCCCTCGGATGGACCTGTTATGCAGAAGAAGACTATGGGATGGGAAGCTTCAAGTGAAAGAATGTACCCTGAAGACGGTGCTCTTAAGGGAGAGATTAAACAACGTCTTAAATTGAAAGATGGAGGACATTACGATGCTGAGGTGAAGACAACTTACAAAGCCAAAAAACCAGTTCAGCTGCCAGGAGCGTACAATGTTAATATTAAACTGGATATCACCTCCCACAACGAGGATTACACTATCGTTGAGCAATATGAAAGAGCTGAAGGGCGGCACTCGACAGGTGGCATGGATGAATTGTATAAG

mCherry 1 synthetic intron 3’ end (ia)

ATGGTCTCAAAGGGTGAAGAAGATAACATGGCAATTATTAAAGAGTTTATGCGTTTCAAGGTGCATATGGAGGGATCTGTCAATGGGCATGAGTTTGAAATTGAAGGTGAAGGAGAAGGCCGACCATATGAGGGAACACAAACCGCAAAACTAAAGGTAACTAAAGGCGGACCATTACCATTCGCCTGGGACATCCTCTCTCCACAGTTCATGTATGGAAGTAAAGCTTATGTTAAACATCCGGCAGATATACCAGATTATTTGAAACTTTCATTCCCGGAGGGTTTTAAGTGGGAACGCGTAATGAATTTTGAAGACGGAGGAGTTGTTACAGTGACGCAAGACTCAAGCCTCCAAGATGGAGAATTTATTTATAAAGTCAAACTTCGAGGAACGAATTTCCCCTCGGATGGACCTGTTATGCAGAAGAAGACTATGGGATGGGAAGCTTCAAGTGAAAGAATGTACCCTGAAGACGGTGCTCTTAAGGGAGAGATTAAACAACGTCTTAAATTGAAAGATGGAGGACATTACGATGCTGAGgtaagtttaaacatatatatactaactaaccctgattatttaaattttcagGTGAAGACAACTTACAAAGCCAAAAAACCAGTTCAGCTGCCAGGAGCGTACAATGTTAATATTAAACTGGATATCACCTCCCACAACGAGGATTACACTATCGTTGAGCAATATGAAAGAGCTGAAGGGCGGCACTCGACAGGTGGCATGGATGAATTGTATAAG

mCherry 1 synthetic intron 5’ end (ic)

ATGGTCTCAAAGGGTGAAGAAGATAACATGGCAATTATTAAAGAGTTTATGCGTTTCAAGGTGCATATGGAGGGATCTGTCAATGGGCATGAGTTTGAAATTGAAGGTGAAGGAGAAGGCCGACCATATGAGGGAACACAAACCGCAAAACTAAAGgtaagtttaaacatgattttactaactaactaatctgatttaaattttcagGTAACTAAAGGCGGACCATTACCATTCGCCTGGGACATCCTCTCTCCACAGTTCATGTATGGAAGTAAAGCTTATGTTAAACATCCGGCAGATATACCAGATTATTTGAAACTTTCATTCCCGGAGGGTTTTAAGTGGGAACGCGTAATGAATTTTGAAGACGGAGGAGTTGTTACAGTGACGCAAGACTCAAGCCTCCAAGATGGAGAATTTATTTATAAAGTCAAACTTCGAGGAACGAATTTCCCCTCGGATGGACCTGTTATGCAGAAGAAGACTATGGGATGGGAAGCTTCAAGTGAAAGAATGTACCCTGAAGACGGTGCTCTTAAGGGAGAGATTAAACAACGTCTTAAATTGAAAGATGGAGGACATTACGATGCTGAGGTGAAGACAACTTACAAAGCCAAAAAACCAGTTCAGCTGCCAGGAGCGTACAATGTTAATATTAAACTGGATATCACCTCCCACAACGAGGATTACACTATCGTTGAGCAATATGAAAGAGCTGAAGGGCGGCACTCGACAGGTGGCATGGATGAATTGTATAAG

mCherry 1 hsp90 intron 5’ end (id)

ATGGTCTCAAAGGGTGAAGAAGATAACATGGCAATTATTAAAGAGTTTATGCGTTTCAAGGTGCATATGGAGGGATCTGTCAATGGGCATGAGTTTGAAATTGAAGGTGAAGGAGAAGGCCGACCATATGAGGGAACACAAACCGCAAAACTAAAGgtttgttttttcgcttctgagtcaattttttaaaatatcggttttagGTAACTAAAGGCGGACCATTACCATTCGCCTGGGACATCCTCTCTCCACAGTTCATGTATGGAAGTAAAGCTTATGTTAAACATCCGGCAGATATACCAGATTATTTGAAACTTTCATTCCCGGAGGGTTTTAAGTGGGAACGCGTAATGAATTTTGAAGACGGAGGAGTTGTTACAGTGACGCAAGACTCAAGCCTCCAAGATGGAGAATTTATTTATAAAGTCAAACTTCGAGGAACGAATTTCCCCTCGGATGGACCTGTTATGCAGAAGAAGACTATGGGATGGGAAGCTTCAAGTGAAAGAATGTACCCTGAAGACGGTGCTCTTAAGGGAGAGATTAAACAACGTCTTAAATTGAAAGATGGAGGACATTACGATGCTGAGGTGAAGACAACTTACAAAGCCAAAAAACCAGTTCAGCTGCCAGGAGCGTACAATGTTAATATTAAACTGGATATCACCTCCCACAACGAGGATTACACTATCGTTGAGCAATATGAAAGAGCTGAAGGGCGGCACTCGACAGGTGGCATGGATGAATTGTATAAG

mEGP no introns

ATGAGTAAAGGAGAAGAACTTTTCACTGGAGTTGTCCCAATTCTTGTTGAATTAGATGGTGATGTTAATGGGCACAAATTTTCTGTCAGTGGAGAGGGTGAAGGTGATGCAACATACGGAAAACTTACCCTTAAATTTATTTGCACTACTGGAAAACTACCTGTTCCATGGCCAACACTTGTCACTACTTTCACTTATGGTGTTCAATGCTTTTCAAGATACCCAGATCATATGAAACgGCATGACTTTTTCAAGAGTGCCATGCCCGAAGGTTATGTACAGGAAAGAACTATATTTTTCAAAGATGACGGGAACTACAAGACACGTGCTGAAGTCAAGTTTGAAGGTGATACCCTTGTTAATAGAATCGAGTTAAAAGGTATTGATTTTAAAGAAGATGGAAACATTCTTGGACACAAATTGGAATACAACTATAACTCACACAATGTATACATCATGGCAGACAAACAAAAGAATGGAATCAAAGTTAACTTCAAAACTAGACACAACATTGAAGATGGAAGCGTTCAACTAGCAGACCATTATCAACAAAATACTCCAATTGGCGATGGCCCTGTCCTTTTACCAGACAACCATTACCTGTCCACACAATCTAAGCTTTCGAAAGATCCCAACGAAAAGAGAGACCACATGGTCCTTCTTGAGTTTGTAACAGCTGCTGGGATTACACATGGCATGGATGAACTATACAAA

mEGFP 3 synthetic introns (ia, ib, ic)

ATGAGTAAAGGAGAAGAACTTTTCACTGGAGTTGTCCCAATTCTTGTTGAATTAGATGGTGATGTTAATGGGCACAAATTTTCTGTCAGTGGAGAGGGTGAAGGTGATGCAACATACGGAAAACTTACCCTTAAATTTATTTGCACTACTGGAAAACTACCTGTTCCATGGgtaagtttaaacatatatatactaactaaccctgattatttaaattttcagCCAACACTTGTCACTACTTTCACTTATGGTGTTCAATGCTTTTCAAGATACCCAGATCATATGAAACgGCATGACTTTTTCAAGAGTGCCATGCCCGAAGGTTATGTACAGGAAAGAACTATATTTTTCAAAGATGACGGGAACTACAAGACACgtaagtttaaacagttcggtactaactaaccatacatatttaaattttcagGTGCTGAAGTCAAGTTTGAAGGTGATACCCTTGTTAATAGAATCGAGTTAAAAGGTATTGATTTTAAAGAAGATGGAAACATTCTTGGACACAAATTGGAATACAACTATAACTCACACAATGTATACATCATGGCAGACAAACAAAAGAATGGAATCAAAGTTgtaagtttaaacatgattttactaactaactaatctgatttaaattttcagAACTTCAAAACTAGACACAACATTGAAGATGGAAGCGTTCAACTAGCAGACCATTATCAACAAAATACTCCAATTGGCGATGGCCCTGTCCTTTTACCAGACAACCATTACCTGTCCACACAATCTAAGCTTTCGAAAGATCCCAACGAAAAGAGAGACCACATGGTCCTTCTTGAGTTTGTAACAGCTGCTGGGATTACACATGGCATGGATGAACTATACAAA
